# Oscillatory correlates of auditory working memory examined with human electrocorticography

**DOI:** 10.1101/2020.06.19.161901

**Authors:** Sukhbinder Kumar, Phillip E. Gander, Joel I. Berger, Alexander J. Billig, Kirill V. Nourski, Hiroyuki Oya, Hiroto Kawasaki, Matthew A. Howard, Timothy D. Griffiths

**Author notes:** **Corresponding authors**: Timothy Griffiths and Sukhbinder Kumar, **Address**: Newcastle University Medical School, Newcastle upon Tyne, Tyne and Wear NE2 4HH, UK, **E-mail** &. These authors contributed equally to the work (Sukhbinder Kumar, Phillip E. Gander, Joel I. Berger). Declaration of Interests: None.

## Abstract

This work examines how sounds are held in auditory working memory (AWM) in humans by examining oscillatory local field potentials (LFPs) in candidate brain regions. Previous fMRI studies by our group demonstrated blood oxygenation level-dependent (BOLD) response increases during maintenance in auditory cortex, inferior frontal cortex and the hippocampus using a paradigm with a delay period greater than 10s. The relationship between such BOLD changes and ensemble activity in different frequency bands is complex, and the long delay period raised the possibility that long-term memory mechanisms were engaged. Here we assessed LFPs in different frequency bands in six subjects with recordings from all candidate brain regions using a paradigm with a short delay period of 3 s. Sustained delay activity was demonstrated in all areas, with different patterns in the different areas. Enhancement in low frequency (delta) power and suppression across higher frequencies (beta/gamma) were demonstrated in primary auditory cortex in medial Heschl’s gyrus (HG) whilst non-primary cortex showed patterns of enhancement and suppression that altered at different levels of the auditory hierarchy from lateral HG to superior- and middle-temporal gyrus. Inferior frontal cortex showed increasing suppression with increasing frequency. The hippocampus and parahippocampal gyrus showed low frequency increases and high frequency decreases in oscillatory activity. The work demonstrates sustained activity patterns that can only be explained by AWM maintenance, with prominent low-frequency increases in medial temporal lobe regions.

## Introduction

In this study we examine the process by which sounds are held in mind over seconds: auditory working memory (AWM). We use the term AWM to refer to the general process of keeping sound objects in mind as opposed to the specific process of phonological working memory to which it is sometimes applied. This general process is relevant to the analysis of auditory scenes in which auditory objects within the scene can remain constant, or come and go. Whilst a number of studies have demonstrated the involvement of early sensory areas in visual working memory (e.g. Pasternak & Greenlee, 2005; Postle, 2006), research examining the role of brain regions in auditory working memory is relatively limited. In particular, although research into neural mechanisms underlying verbal working memory is growing (e.g. Fiebach, et al., 2006, 2006; Ivanova, et al., 2018), the neural substrates of AWM in general are not well established.

Previous research into AWM used functional magnetic resonance imaging (fMRI), magnetoencephalography or positron emission tomography. With respect to sensory cortex, data from these studies are conflicting; whilst the majority of studies found increased activity in auditory cortex during the delay period (Gaab, et al., 2003; Grimault, et al., 2009; Kumar, et al., 2016; Linke & Cusack, 2015), others showed suppression of activity (Linke, et al., 2011; Zatorre, et al., 1994). It should be noted that some of the original studies did not allow for separation of the delay period from encoding or retrieval periods of working memory, a factor which could explain discrepancies. With respect to a broader network involved in AWM maintenance, previous functional imaging has implicated the inferior frontal lobe and the hippocampus in maintenance of AWM (Kumar, et al., 2016). However, that fMRI experiment used a long delay period to overcome the slow hemodynamic response (in order to demonstrate delay period activity ‘uncontaminated’ by encoding or retrieval activity) and may have engaged long-term memory mechanisms.

In this study we carried out LFP recordings from areas previously implicated in AWM maintenance using BOLD measurement. This allows examination of neural activity that is spatially and temporally precise in order to assess patterns of activity across frequency bands. The relationship between such LFP patterns and BOLD changes is complex. For example, studies of sensory cortex suggest a positive correlation between high-frequency LFP oscillations and BOLD (Mukamel, et al., 2005; Oya, et al., 2018) and negative correlation with lower (delta, theta) frequency bands, but in hippocampus this relationship does not seem to hold as a positive correlation between theta/delta frequency band and BOLD exists (Figure 1 in Ekstrom, 2010). More fundamentally, the technique is a direct test of whether explicit sustained activity during maintenance occur in parts of the suggested brain network. This is not a given, based on considerable recent interest in ‘activity silent’ maintenance of working memory (Reynolds, et al., 2009; for a recent review, see Sreenivasan & D’Esposito, 2019). Working with LFPs also allows us to use a paradigm with a short delay period during which activity can only be interpreted on the basis of working memory.

**Figure 1.**
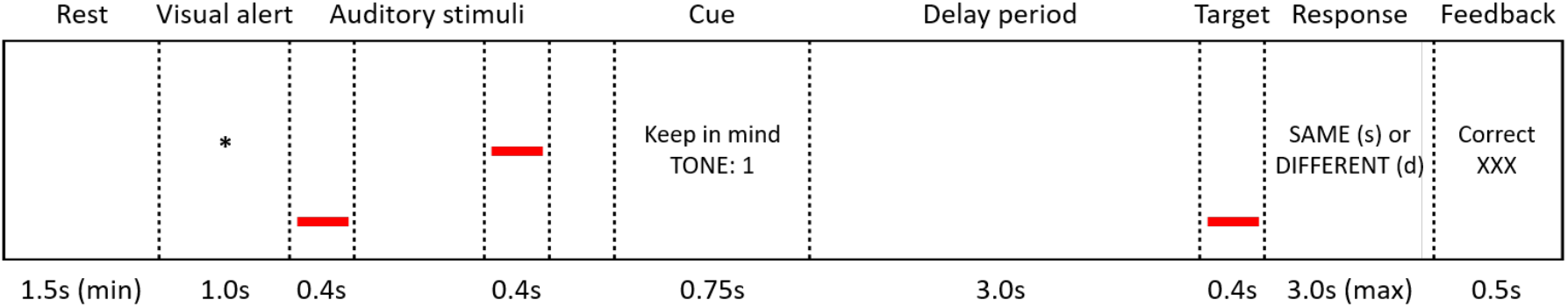
Paradigm for working memory task. Subjects were cued to maintain a particular tone for a period of three seconds following the presentation of two different tones. They were then asked to compare the maintained tone with a target tone, and given feedback as to whether they were correct, followed by a rest period.

## Methods

### Subjects

Six neurosurgical patients were studied (all six male, right-handed with left language dominance, age 17 – 39 years old); five were within the low to high average intellect range with normal working memory, while one patient (L275) was cognitively impaired and deficient in working memory on neuropsychological assessment. They had medically refractory epilepsy for which electrocorticography was required to identify seizure foci prior to potential resection surgery. Electrode contacts in the region of the seizure focus were not included in further analysis. Research protocols were examined and approved by the University of Iowa Institutional Review Board.

### Electrodes and recording setup

Recordings were made from subjects in an electromagnetically shielded facility. Intracranial data were recorded from the following electrode arrays implanted on the right ride (N = 3, as indicated by the letter prefix ‘R’ in the subject code) and the left (N = 3, prefix code ‘L’): an 8-contact depth electrode (5 mm center-to-center distance) placed along the axis of Heschl’s gyrus (Howard, et al., 1996; Nagahama, et al., 2018; Reddy, et al., 2010); a 96-contact grid (5 mm center-to-center distance) covering a large area of the temporal lobe (and in one subject portions of the inferior frontal gyrus; IFG); multi-contact grids (16 or 32 contacts) overlying inferior frontal cortex; and a strip electrode (12 contacts) placed medially along the anterior ventral portion of the temporal lobe. All electrodes (Ad-Tech Medical Instrument, Racine, WI and PMT Corporation, Chanhassen, MN) were digitized using a Tucker Davis Technologies (Alachua, FL) RZ2 system and referenced to a subgaleal electrode. Data were acquired using a 0.7-800 Hz bandpass filter (sampled at 2034 Hz) at 16-bit resolution. Electrode placement was selected based purely on clinical considerations for seizure monitoring. Regions of interest (ROIs) for the purposes of analysis were guided by our previous construction of an auditory working memory network (Kumar, et al., 2016), but with greater spatial precision than was available when using fMRI. These included posteromedial Heschl’s gyrus (HGpm), anterolateral Heschl’s gyrus (HGal), superior temporal gyrus (STG), middle temporal gyrus (MTG), inferior frontal gyrus (IFG), hippocampus (HC) and parahippocampal gyrus (PHG).

### Stimuli and procedure

Auditory stimuli were delivered diotically using Etymotic ER4B earphones with custom-made earmolds. Sounds were set at a level that was confirmed to be comfortable for each subject. The experimental procedure was based on that used in Kumar et al. (2016) shown in Figure 1. For each trial, subjects were presented with a visual alert which indicated the start of a trial. Following a one second delay, two tones (400 ms duration, 500 ms for subject L275, gated with a cosine window of 10ms at the onset and offset), selected randomly, one from each of the two sets consisting of 8 tones, logarithmically sampled from a low frequency (between 300-570 Hz) or high frequency (between 2000-2800 Hz) range. A silent inter-stimulus interval of 750 ms (1 s for L275) separated the two tones. After another delay of 250 ms (500 ms for L275), a visual cue was presented for 750 ms (1.5 s for L275), instructing the subject to remember either the first or the second tone. This was followed by a 3000 ms period during which the subjects were required to remember the tone, referred to as the “delay period”. Finally, subjects were presented with a target tone which on half of the trials was the same tone as they were asked to remember, and on the other half of trials was either + or − 20% of the cued tone frequency. They were then visually instructed to respond on the keyboard whether this target tone was the same or different to the tone they had been asked to remember. They were given feedback as to whether they were correct or not, which was followed by a short rest period (4500-5500 ms) prior to the start of the subsequent trial. The total trial duration was 14.25-16.7 seconds. Subjects completed up to 160 trials in total.

### Imaging

A high-resolution T1-weighted structural MRI of the brain was acquired for each subject before and after electrode implantation. Images were acquired from a 3T Siemens TIM Trio scanner with a 12-channel head coil (MPRAGE: 0.78 × 0.78 mm, slice thickness 1.0 mm, TR = 2.530 s, TE = 3.520 ms, average of two). To determine the location of recording contacts on the preoperative structural MRI, these images were coregistered to post-implantation structural MRIs using a 3D linear registration algorithm (Functional MRI of the Brain Linear Image Registration Tool; Jenkinson, etal., 2012) and custom Matlab v.9.0 (MathWorks, Natick, MA) scripts using guidance from computed tomography (CT) scans (in-plane resolution 0.51 × 0.51 mm, slice thickness 1.0 mm). When possible, results were also compared with intraoperative photographs to ensure reconstruction accuracy. To be included, all contacts were confirmed to be in grey matter within each ROI. Each individual’s brain was coregistered to the ICBM152 template (Fonov, et al., 2011) to estimate contact location in MNI template space.

### Data processing and analysis

Data were analysed offline using custom-written Matlab scripts and the EEGLab toolbox (Delorme & Makeig, 2004). In order to probe successful working memory processes, only trials where the subject responded correctly were included. Data were downsampled to 500 Hz and trials were epoched, with times ranging from 2000 ms prior to the initial visual alert to 11000 ms after. Line noise was removed by implementing the demodulated band transform (Kovach & Gander, 2016). Data were further denoised by removing the first principal component from the singular value decomposition of the highpass-filtered data (> 160 Hz).

Morlet-based wavelet analyses were performed to examine changes in frequency-specific activity over time. Trials were first rejected using automated artifact rejection, based on kurtosis and large common amplitudes across electrodes (Delorme, et al., 2007). Remaining data were processed in sliding windows, using a minimum of 3 cycles per window for the lowest frequency, increasing linearly with increasing frequency. Wavelet analyses were calculated between 2-150 Hz. Time-frequency data were normalized to the average power across trials. Data were permuted across time and trials, and a significance threshold of 0.01 (two-tailed) was set for the purposes of displaying the data, relative to these permuted values, and responses for the whole trial were compared to a pre-alert baseline period (−1000 to −100 ms). Band power during the delay period was calculated by taking the mean power across time and frequency within canonical frequency bands, omitting the first and last 100 ms to account for the width of the Gaussian window. These were delta (2-4Hz), theta (4-8 Hz), alpha (8-13 Hz), beta (13-30 Hz), low gamma (30-70 Hz) and high gamma (70-150 Hz). To reduce the dimensionality of the data, time-frequency data were selected from the electrode exhibiting the strongest absolute response (i.e. regardless of directionality) within any frequency band, which was confirmed to be representative of surrounding electrodes, for each ROI.

To further examine the dynamics of the oscillatory activity, cluster-based permutation tests (Maris & Oostenveld, 2007) were run, comparing the power in each canonical frequency range during the delay period to the average power during the baseline period, for each ROI. This involved running two-tailed, paired *t*-tests at every timepoint. Data were clustered on the basis of temporal adjacency, with a pre-designated threshold of *p* < 0.05 set for determining the *t*-values required. The largest summed *t*-value cluster was used as the test statistic. Data were then shuffled between baseline and the delay period and this method was repeated for all possible permutations, to create a distribution of the largest summed *t*-value clusters. Summed *t*-value clusters from the actual data that fell outside the 95% confidence interval of this distribution were considered significant.

## Results

Percentage accuracy of the subjects in the task varied from 67% to 99% (Figure 2). All subjects performed above chance level (chi-square test, p < 0.05).

**Figure 2:**
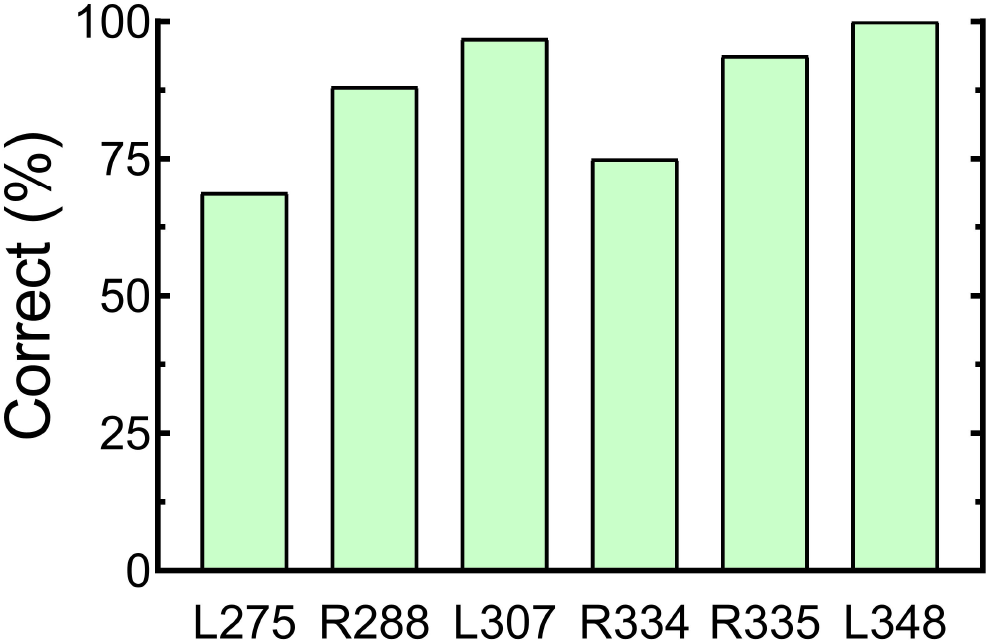
Behavioral performance of all subjects in the task

Example event-related spectral perturbations (ERSPs) from one subject are shown in Figure 3. The ERSPs for remaining subjects are included as supplementary figures (Supplementary Figures 1-5). Activity during the delay period is described below for each ROI. Data across subjects are summarized as median changes in power (dB) relative to baseline. These data are shown in Figure 4.

**Figure 3.**
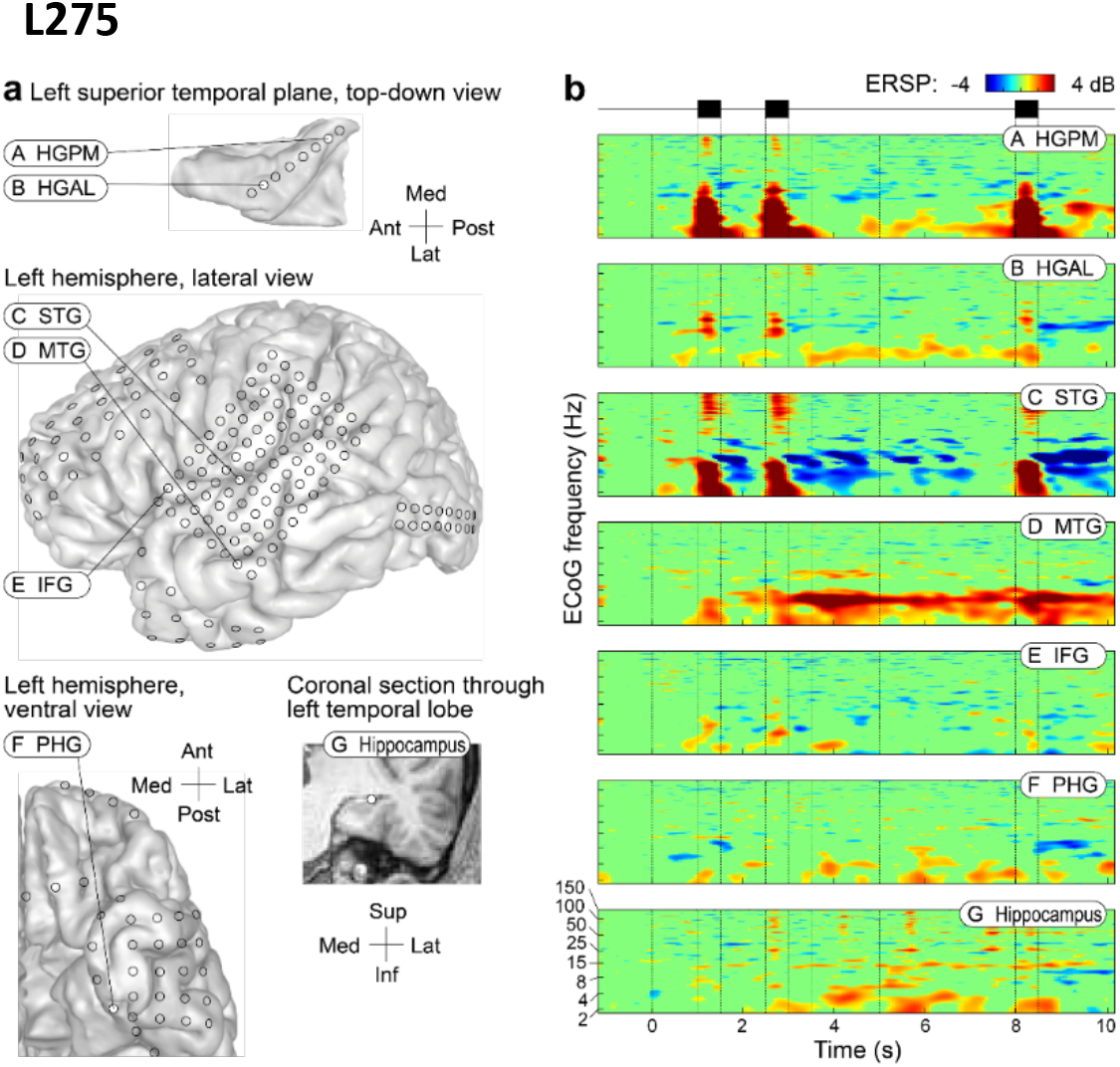
Example intracranial data from a single patient (L275). **a.** Location of implanted electrodes, with an 8-channel depth electrode along the axis of HG, a single 96-channel grid over STG, MTG and IFG, a strip electrode overlying the PHG and a depth electrode into HC. **b.** Corresponding event-related spectral perturbations (ERSPs) recorded from the ROIs during the working memory task.

**Figure 4.**
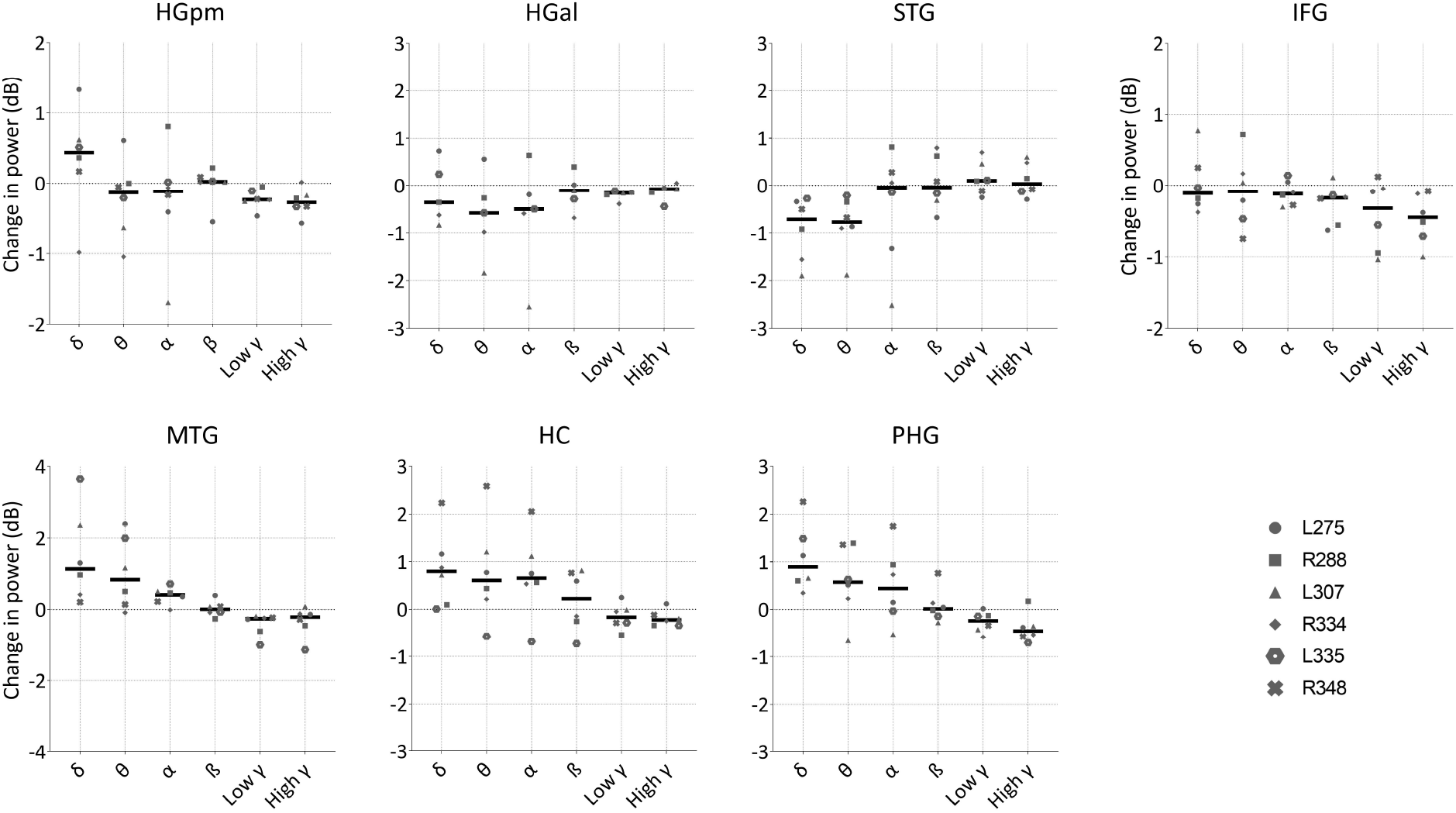
Change in power during the delay period, for each canonical frequency band and each ROI, plotted across subjects (*n* = 6). Thick coloured lines represent median across subjects. Grey markers indicate individual subject data. Axes are scaled to the maximum value for each ROI.

### HGpm

During the delay period, increases in delta band power were evident in HGpm in five of the six subjects, with a median value of 0.44 dB. Activity in other frequency bands was either unchanged across subjects (beta = 0.02 dB) or decreased to varying degrees (theta = −0.13; alpha = −0.12 dB; low gamma −0.24 dB; high gamma −0.27 dB).

Data were also plotted over the whole delay period, to examine the temporal dynamics of oscillatory activity (Figure 5). With the exception of the one subject who exhibited a decrease in activity, the increases in delta band activity were not restricted to a particular period but were distributed across the whole epoch after the first 500 ms. In contast, a transient decrease in theta band activity occurred that was restricted to the first ~750 ms of the delay period in all but one subject (Figure 5B), whilst decreases in low and high gamma and, to a lesser extent, alpha band, were sustained throughout the period (Figure 5C, 5E, and 5F). However, when examined on a moment-by-moment basis, only the decreases in low and high gamma during the delay period were significantly different from baseline (cluster-based permutation test; *p* < 0.05). Note that the sharp increase of power in delta/theta bands towards the end of the delay period is due to the long analysis window for these bands, which overlaps with the upcoming sound period.

**Figure 5.**
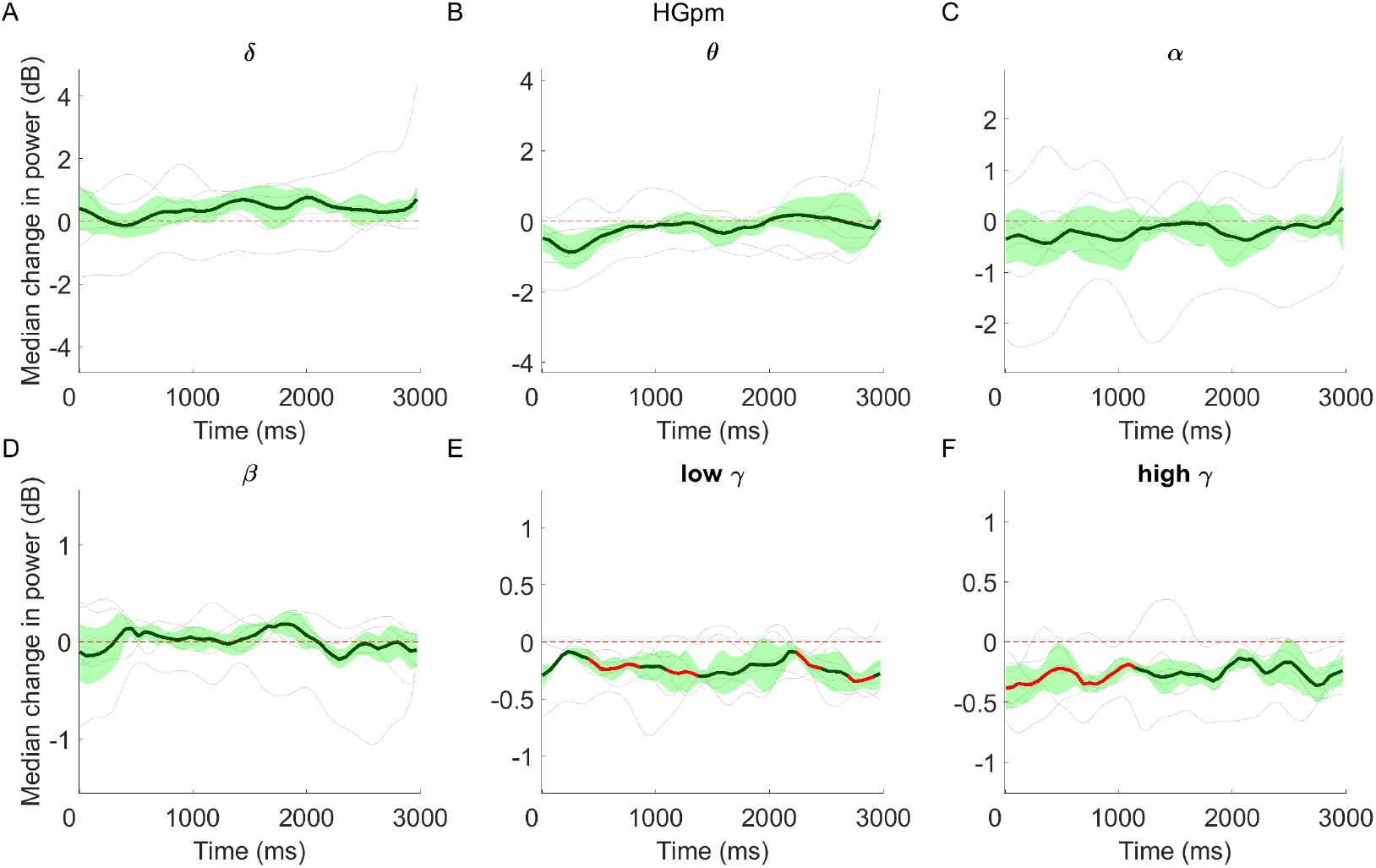
Median change in power across subjects, relative to baseline, plotted across the entire delay period for HGpm. Time referenced to start of delay period. Green shading indicates median absolute deviation (MAD), light grey lines show individual subject data, whilst red areas highlight periods of significance, determined using cluster-based permutation tests with an alpha level of 0.05.

### HGal

Median activity in HGal was decreased in all frequency bands: delta (−0.36 dB), theta (−0.58 dB), alpha (−0.50 dB), beta (−0.11 dB), low gamma (−0.16 dB) and high gamma (−0.08 dB). Analysis of activity over time revealed that decreases in delta through alpha band activity were generally restricted to the first ~750 ms of the delay period (in all but one subject). As with HGpm, decreases in low gamma and high gamma were evident throughout the whole period, although these were only significant across subjects for low gamma band (Supplementary Figure 6).

### STG

Large decreases in delta (−0.71 dB) and theta activity (−0.78 dB) were evident in STG during the delay period. Median alpha, beta and high gamma band activity was relatively unchanged (−0.04 dB,−0.04 dB and 0.03 dB respectively), although this was variable across subjects. Slight increases occurred in low gamma (0.1 dB). When examined over time across subjects, significant decreases in delta and theta were evident during the first 1000 ms of the delay period (cluster-based permutation test, *p* < 0.05; Supplementary Figure 7).

### MTG

Clear increases in delta (1.10 dB) and theta activity (0.83 dB) were present in MTG during the delay period. When examined across subjects and time, these changes were sustained throughout the whole period but only reached significance sporadically (Supplementary Figure 8). More moderate increases were also evident in alpha (0.41 dB), whilst low and high gamma were decreased (−0.26 dB and −0.21 dB, respectively), and beta band activity was unchanged (0dB). Decreases in low gamma were significant for the first ~600 ms but power in this band remained suppressed for the duration of the delay period.

### IFG

Median band power in IFG decreased with increasing frequency. Slight decreases were evident in delta (−0.11 dB), theta (−0.08 dB) and alpha (−0.12 dB) bands, and larger decreases were present in beta (−0.17 dB), low gamma (−0.32 dB) and high gamma bands (−0.45 dB). Although delta band activity was unchanged when averaging band power across the whole delay period, there was a significant decrease evident in the final ~300 ms (Figure 6A). Decreases in beta were sustained for the majority of the delay period, although these showed an upward trend towards the final 750 ms (Figure 5D). Gamma band activity was suppressed throughout the epoch, but only reached significance across subjects for high gamma in the second half of the delay period (Figure 6E-6F).

**Figure 6.**
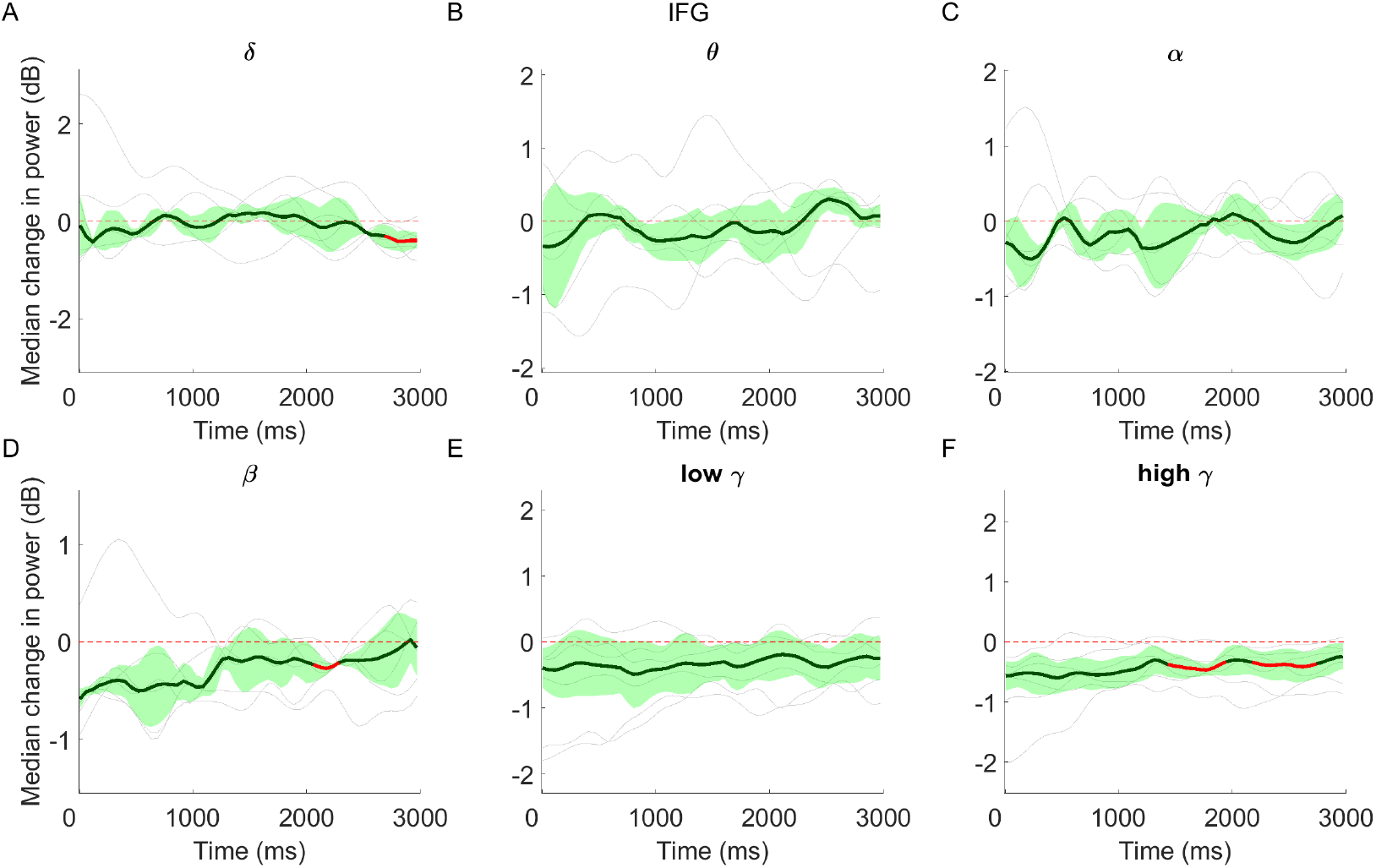
Median change (± MAD) in power relative to baseline, across subjects and over time, in IFG. Red areas highlight periods of significance. Single subject data are overlaid (light grey lines).

### HC

Across subjects, clear increases in hippocampus oscillatory activity were evident in delta (0.80 dB), theta (0.61 dB), alpha (0.66 dB) and - to a lesser extent - beta band (0.22 dB). Median activity in low and high gamma bands decreased (−0.17 dB and −0.22 dB, respectively). Temporal presentation of the data revealed that the increases in delta band activity were sustained throughout the delay period whilst other bands generally showed a build-up within the first 1000 ms and then remained sustained (Figure 7A). Due to variability across subjects, these increases did not generally reach significance, other than for a short section of the delay period for delta activity. Low gamma activity was significantly reduced across subjects during the first ~500 ms of the delay period, whilst significant decreases in high gamma activity were periodic and not sustained for the whole period.

**Figure 7.**
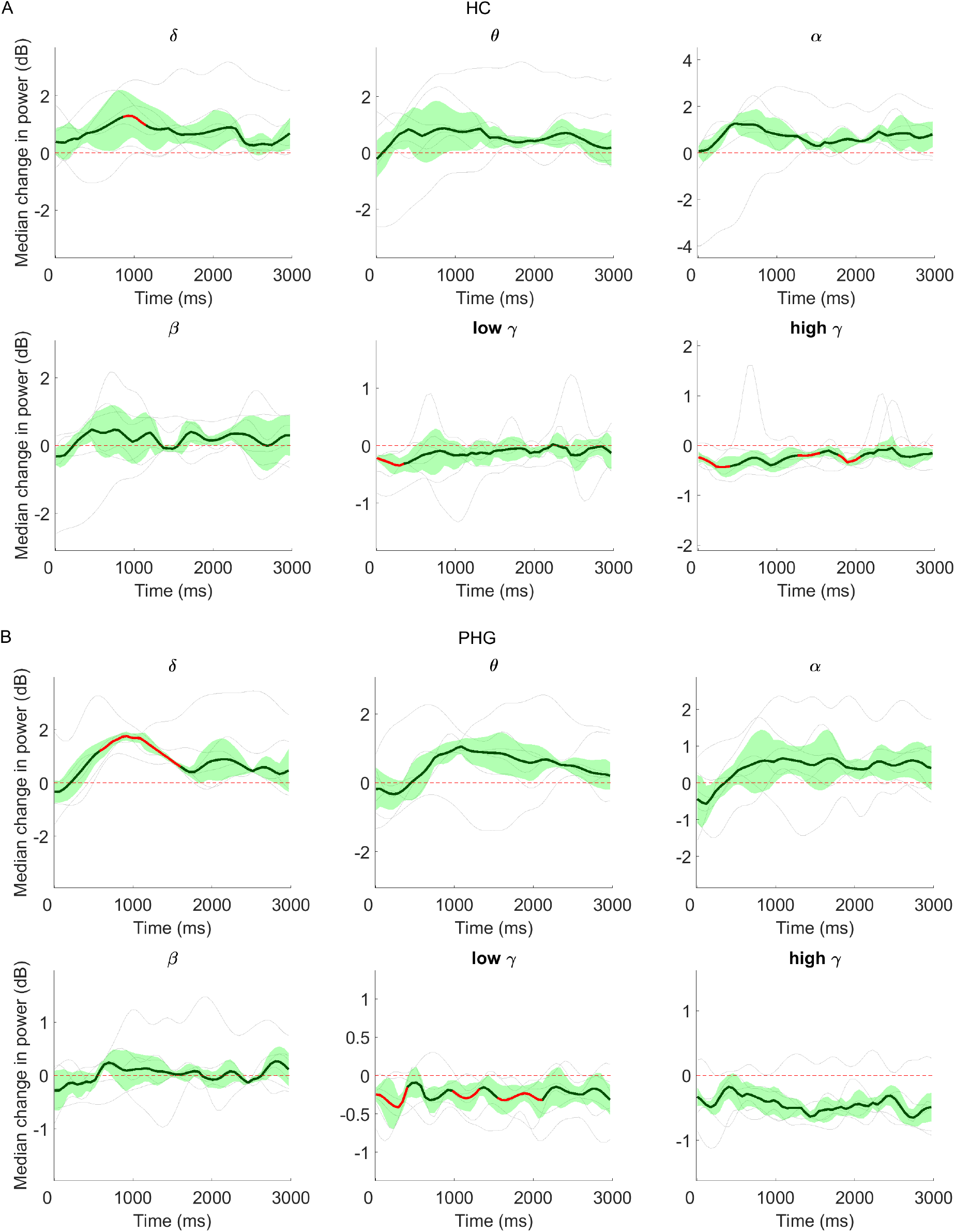
Median change (± MAD) in power across subjects, relative to baseline, plotted across the entire delay period for HC **(A)** and PHG **(B)**. Red areas highlight periods of significance. Single subject data are overlaid (light grey lines).

### PHG

Clear increases were evident in delta band activity in all subjects (0.90 dB). Similar to HC, increases were also evident in theta (0.57 dB) and alpha bands (0.44 dB), whilst decreases were present in low gamma (−0.24 dB) and high gamma (−0.46 dB). Median beta band activity was relatively unchanged (0.01 dB). Increases in low frequency activity showed a build-up within the first ~750 ms and were then evident for the remainder of the delay period when examined over time (Figure 7B). Due to some variability across subjects, these increases were significant only from 500 to 1500 ms for delta band activity. Decreases in both low and high gamma band activity were generally sustained for the entire delay period.

## Discussion

We have demonstrated activity during auditory working memory maintenance in all three candidate areas: auditory cortex, inferior frontal cortex and medial temporal lobe. Delta power increases and suppression across higher frequencies were found in primary auditory cortex in medial Heschl’s gyrus (HG) whilst non-primary cortex showed patterns of activation and suppression that altered at different levels of the auditory hierarchy from lateral HG to superior and middle temporal gyrus. Inferior frontal cortex showed increasing suppression with increasing frequency. The hippocampus and parahippocampal gyrus showed low-frequency increases and high-frequency decreases in oscillatory activity.

### Delay activity in primary auditory cortex

Above-baseline delta power was previously identified in auditory cortex while subjects kept tone frequencies in mind for a brief interval (0.5-3s; Herbst & Obleser, 2019). While that scalp EEG study could not distinguish between auditory fields, the delta power increase we report is specific to primary cortex in posteromedial Heschl’s gyrus. Decreases in power for other low frequencies (theta and alpha) and high frequencies (gamma and high gamma) were observed in the primary auditory cortex. In our previous study, we demonstrated an increase in BOLD response in the same region during a longer delay period (Kumar, et al., 2016). This can be reconciled with our current findings by recalling that increase in BOLD signal can be either due to decrease in low frequencies (delta, theta, alpha) and/or increase in gamma/high gamma power. Also, in terms of predicting a BOLD signal, the different frequency bands are not weighted equally. For example, theta and alpha frequencies have a higher (negative) weightage than delta band and (positive) weightage of gamma/high gamma is lower than (negative) weightage of theta/alpha bands (Mukamel, et al., 2005; Zumer et al., 2010). There is also some uncertainty regarding the lowest frequency up to which BOLD has a negative correlation. While Mukamel, et al., (2005) show this to be beta band, other studies e.g. Zumer et al. (2010) show negative correlation of BOLD extends up to lower gamma band (~50Hz). Therefore, decreases in theta/alpha (and possibly gamma) band combined with their stronger weightings could explain the increase in BOLD response observed in our previous work (Kumar, et al., 2016).

Recent theoretical and empirical work points to short-term updating of synaptic weights in sensory cortex as one mechanism for working memory maintenance (Masse, et al., 2019; Miller, et al., 2018; Sreenivasan & D’Esposito, 2019; Wolff, et al., 2017; Wolff, et al., 2020). Such effects are described as activity-silent: they are not directly measurable as changes in neural firing or local field potentials but may be inferred through their impact on neural responses to brief task-irrelevant impulse stimuli presented during the maintenance period (Wolff, et al., 2017; Wolff, et al., 2020). While this approach holds promise, our data show that keeping the frequency of a tone in mind can in fact generate sustained oscillatory signatures in auditory cortex and beyond. Further work is required to establish the relationship between the LFP delta increases that are seen here, neuronal activity, and measures of the synaptic ‘trace’ that might maintain AWM.

### Delay activity in higher-order lateral temporal cortex

Non-primary auditory cortex in HGal and STG did not show the same delta power increase as HGpm; indeed power in these regions was more suppressed across low-frequencies (delta-alpha). This distinction between oscillatory signatures in primary and non-primary auditory fields is similar to that observed during active listening to speech (Billig, et al., 2019). In our previous fMRI study (Kumar, et al., 2016), we showed a greater increase in BOLD signal (compared to primary auditory cortex) as we moved up the hierarchy to lateral HG and planum temporale. This is consistent (as per the discussion above) with a stronger decrease in power of all low frequencies (delta, theta, alpha) in HGal and with decreased power of delta/theta band in STG. There is not much change in the power of gamma/high gamma band in these regions.

Moving up the hierarchy, a contrasting and robust above-baseline increase in delta and theta power was observed in MTG. Interestingly, this oscillatory pattern is more similar to that in the medial temporal lobe (PHG/HC) than in hierarchically earlier auditory sites. We had no prior expectations of activity in this region. MTG is known to represent high-level sound categories during imagery (Linke & Cusack, 2015) and to be involved in semantic processing more generally (Davey, et al., 2016; Hickok & Poeppel, 2004). Despite the simple nature of the pure tone stimuli, it is possible that subjects tried to apply semantic labels in performing the task.

### Delay activity in inferior frontal gyrus

Above-baseline BOLD activity and PET regional cerebral blood flow have been reported in inferior frontal gyrus during the maintenance and comparison of pure tones (Zatorre, et al., 1994; Kumar et al., 2016) and amplitude modulation rate (Uluc, et al., 2018). The present study demonstrates suppression of LFP activity in inferior frontal cortex that is greatest at high frequency and a less pronounced decrease in power of lower frequencies. If we use the same rationale as used for auditory cortex in the previous sections, then the oscillatory pattern of IFG would predict a decrease rather than an increase in BOLD signal, which is inconsistent with our previous fMRI work. However, two points need to be kept in mind before this conclusion is drawn. First, the relation between oscillatory power and BOLD in the IFG may not be same as in sensory cortex. For example, while beta band power (20-30Hz) is positively correlated with BOLD in the auditory cortex (Mukamel, et al., 2005), power in the same frequency band is highly negatively correlated with BOLD signal in frontal regions (Hanslmayr, et al., 2011). Second, IFG coverage in the current study does not the overlap with the region identified in the fMRI study, which is much more ventral (frontal operculum bordering on anterior insula).

### Delay activity in the medial temporal lobe

Although our previous fMRI work identified hippocampal activity in support of auditory working memory (Kumar et al., 2016), the prolonged delay period in that study may have engaged a long-term or episodic hippocampal store (Scoville & Milner, 1957; Squire, 1992). In the current experiment, the temporal resolution of electrophysiological recordings allowed us to reduce the interval to 3 seconds and still dissociate encoding from delay activity. The pronounced power increases observed in (para)hippocampal delta-theta here are consistent with the fMRI findings (Kumar et al., 2016), given the positive correlation between low frequency power and the BOLD signal in hippocampus (Ekstrom, 2010). They are also reminiscent of hippocampal signatures of successful spatial and episodic encoding of memories tested over longer timeframes (Cornwell, et al., 2008; Lega, et al., 2012). Theta oscillations and the coupling of their phase with firing and high frequency activity have also been associated with visual and phonological working memory beyond the medial temporal lobe (Raghavachari, et al., 2001, Canolty, et al., 2006).

While some previous work emphasises a role for the medial temporal lobe in working memory when maintaining complex, conjunctive, or novel features (reviewed in Sreenivasan & D’Esposito, 2019), the tone frequencies we manipulated form a fundamental sound dimension. Rodent work has shown tuning for this feature in hippocampus and entorhinal cortex when it is task relevant (Aronov, et al., 2017) although we do not know whether this is the case in humans. On the other hand, cells in hippocampus that fire in a fixed sequence to span a delay period have been identified both in rodents (Pastalkova, et al., 2008; MacDonald, et al., 2013) and humans (Umbach, et al., 2020). Their ability to support working memory may depend on input from adjoining parahippocampal fields (Suh, et al., 2011; Robinson, et al., 2017).

### Conclusions and outlook

The work demonstrates patterns of activity that can only be explained by AWM maintenance, that are sustained during a delay interval, with prominent low-frequency increases in medial temporal lobe regions. Establishing the relationship between this medial temporal activity and that in areas more routinely associated with AWM, including inferior frontal gyrus and auditory cortex, will be an important next step. The hippocampus and inferior frontal gyrus may help keep sensory traces active in auditory cortex via ongoing top-down signaling, or alternatively maintain independent representations at different degrees of abstraction from the stimulus (Kumar et al., 2016; Huang, et al., 2016). Future studies that draw on effective connectivity and pattern similarity analyses will be key to distinguishing between these and other accounts.

## Acknowledgements

This work was supported by Wellcome Trust grant (WT106964MA), NIH grant (DC004290) and Hoover Fund. This work was conducted on an MRI instrument funded by 1S10OD025025-01.

## References

Aronov, D., Nevers, R., & Tank, D. W. (2017). Mapping of a non-spatial dimension by the hippocampal-entorhinal circuit. Nature, 543, 719–722.

Billig, A. J., Herrmann, B., Rhone, A. E., Gander, P. E., Nourski, K. V., Snoad, B. F., Kovach, C. K., Kawasaki, H., Howard, M. A., 3rd, & Johnsrude, I. S. (2019). A Sound-Sensitive Source of Alpha Oscillations in Human Non-Primary Auditory Cortex. J Neurosci, 39, 8679–8689.

Canolty, R. T., Edwards, E., Dalal, S. S., Soltani, M., Nagarajan, S. S., Kirsch, H. E., Berger, M. S., Barbaro, N. M., & Knight, R. T. (2006). High gamma power is phase-locked to theta oscillations in human neocortex. Science, 313, 1626–1628.

Cornwell, B. R., Johnson, L. L., Holroyd, T., Carver, F. W., & Grillon, C. (2008). Human hippocampal and parahippocampal theta during goal-directed spatial navigation predicts performance on a virtual Morris water maze. J Neurosci, 28, 5983–5990.

Davey, J., Thompson, H. E., Hallam, G., Karapanagiotidis, T., Murphy, C., De Caso, I., Krieger-Redwood, K., Bernhardt, B. C., Smallwood, J., & Jefferies, E. (2016). Exploring the role of the posterior middle temporal gyrus in semantic cognition: Integration of anterior temporal lobe with executive processes. Neuroimage, 137, 165–177.

Delorme, A., & Makeig, S. (2004). EEGLAB: an open source toolbox for analysis of single-trial EEG dynamics including independent component analysis. J Neurosci Methods, 134, 9–21.

Delorme, A., Sejnowski, T., & Makeig, S. (2007). Enhanced detection of artifacts in EEG data using higher-order statistics and independent component analysis. Neuroimage, 34, 1443–1449.

Ekstrom, A. (2010). How and when the fMRI BOLD signal relates to underlying neural activity: the danger in dissociation. Brain Res Rev, 62, 233–244.

Fiebach, C. J., Rissman, J., & D’Esposito, M. (2006). Modulation of inferotemporal cortex activation during verbal working memory maintenance. Neuron, 51, 251–261.

Fonov, V., Evans, A. C., Botteron, K., Almli, C. R., McKinstry, R. C., Collins, D. L., & Brain Development Cooperative G. (2011). Unbiased average age-appropriate atlases for pediatric studies. Neuroimage, 54, 313–327.

Gaab, N., Gaser, C., Zaehle, T., Jancke, L., & Schlaug, G. (2003). Functional anatomy of pitch memory--an fMRI study with sparse temporal sampling. Neuroimage, 19, 1417–1426.

Grimault, S., Lefebvre, C., Vachon, F., Peretz, I., Zatorre, R., Robitaille, N., & Jolicoeur, P. (2009). Load-dependent brain activity related to acoustic short-term memory for pitch: magnetoencephalography and fMRI. Ann N Y Acad Sci, 1169, 273–277.

Hanslmayr, S., Volberg, G., Wimber, M., Raabe, M., Greenlee, M. W., & Bauml, K. H. (2011). The relationship between brain oscillations and BOLD signal during memory formation: a combined EEG-fMRI study. J Neurosci, 31, 15674–15680.

Herbst, S. K., & Obleser, J. (2019). Implicit temporal predictability enhances pitch discrimination sensitivity and biases the phase of delta oscillations in auditory cortex. Neuroimage, 203, 116198.

Hickok, G., & Poeppel, D. (2004). Dorsal and ventral streams: a framework for understanding aspects of the functional anatomy of language. Cognition, 92, 67–99.

Howard, M. A., 3rd, Volkov, I. O., Granner, M. A., Damasio, H. M., Ollendieck, M. C., & Bakken, H. E. (1996). A hybrid clinical-research depth electrode for acute and chronic in vivo microelectrode recording of human brain neurons. Technical note. J Neurosurg, 84, 129–132.

Huang, X., Chen, X., Yan, N., Jones, J. A., Wang, E. Q., Chen, L., Guo, Z., Li, W., Liu, P., & Liu, H. (2016). The impact of parkinson’s disease on the cortical mechanisms that support auditory-motor integration for voice control. Hum Brain Mapp, 37, 4248–4261.

Ivanova, M. V., Dragoy, O., Kuptsova, S. V., Yu Akinina, S., Petrushevskii, A. G., Fedina, O. N., Turken, A., Shklovsky, V. M., & Dronkers, N. F. (2018). Neural mechanisms of two different verbal working memory tasks: A VLSM study. Neuropsychologia, 115, 25–41.

Jenkinson, M., Beckmann, C. F., Behrens, T. E., Woolrich, M. W., & Smith, S. M. (2012). Fsl. Neuroimage, 62, 782–790.

Kovach, C. K., & Gander, P. E. (2016). The demodulated band transform. J Neurosci Methods, 261, 135–154.

Kumar, S., Joseph, S., Gander, P. E., Barascud, N., Halpern, A. R., & Griffiths, T. D. (2016). A Brain System for Auditory Working Memory. J Neurosci, 36, 4492–4505.

Lega, B. C., Jacobs, J., & Kahana, M. (2012). Human hippocampal theta oscillations and the formation of episodic memories. Hippocampus, 22, 748–761.

Linke, A. C., & Cusack, R. (2015). Flexible information coding in human auditory cortex during perception, imagery, and STM of complex sounds. J Cogn Neurosci, 27, 1322–1333.

Linke, A. C., Vicente-Grabovetsky, A., & Cusack, R. (2011). Stimulus-specific suppression preserves information in auditory short-term memory. Proc Natl Acad Sci U S A, 108, 12961–12966.

MacDonald, C. J., Carrow, S., Place, R., & Eichenbaum, H. (2013). Distinct hippocampal time cell sequences represent odor memories in immobilized rats. J Neurosci, 33, 14607–14616.

Maris, E., & Oostenveld, R. (2007). Nonparametric statistical testing of EEG- and MEG-data. J Neurosci Methods, 164, 177–190.

Masse, N. Y., Yang, G. R., Song, H. F., Wang, X. J., & Freedman, D. J. (2019). Circuit mechanisms for the maintenance and manipulation of information in working memory. Nat Neurosci, 22, 1159–1167.

Miller, E. K., Lundqvist, M., & Bastos, A. M. (2018). Working Memory 2.0. Neuron, 100, 463–475.

Mukamel, R., Gelbard, H., Arieli, A., Hasson, U., Fried, I., & Malach, R. (2005). Coupling between neuronal firing, field potentials, and FMRI in human auditory cortex. Science, 309, 951–954.

Nagahama, Y., Kovach, C. K., Ciliberto, M., Joshi, C., Rhone, A. E., Vesole, A., Gander, P. E., Nourski, K. V., Oya, H., Howard, M. A., Kawasaki, H., & Dlouhy, B. J. (2018). Localization of musicogenic epilepsy to Heschl’s gyrus and superior temporal plane: case report. J Neurosurg, 129, 157–164.

Oya, H., Gander, P. E., Petkov, C. I., Adolphs, R., Nourski, K. V., Kawasaki, H., Howard, M. A., & Griffiths, T. D. (2018). Neural phase locking predicts BOLD response in human auditory cortex. Neuroimage, 169, 286–301.

Pastalkova, E., Itskov, V., Amarasingham, A., & Buzsaki, G. (2008). Internally generated cell assembly sequences in the rat hippocampus. Science, 321, 1322–1327.

Pasternak, T., & Greenlee, M. W. (2005). Working memory in primate sensory systems. Nat Rev Neurosci, 6, 97–107.

Postle, B. R. (2006). Working memory as an emergent property of the mind and brain. Neuroscience, 139, 23–38.

Raghavachari, S., Kahana, M. J., Rizzuto, D. S., Caplan, J. B., Kirschen, M. P., Bourgeois, B., Madsen, J. R., & Lisman, J. E. (2001). Gating of human theta oscillations by a working memory task. J Neurosci, 21, 3175–3183.

Reddy, C. G., Dahdaleh, N. S., Albert, G., Chen, F., Hansen, D., Nourski, K., Kawasaki, H., Oya, H., & Howard, M. A., 3rd. (2010). A method for placing Heschl gyrus depth electrodes. J Neurosurg, 112, 1301–1307.

Reynolds, J. R., West, R., & Braver, T. (2009). Distinct neural circuits support transient and sustained processes in prospective memory and working memory. Cereb Cortex, 19, 1208–1221.

Robinson, N. T. M., Priestley, J. B., Rueckemann, J. W., Garcia, A. D., Smeglin, V. A., Marino, F. A., & Eichenbaum, H. (2017). Medial Entorhinal Cortex Selectively Supports Temporal Coding by Hippocampal Neurons. Neuron, 94, 677–688 e676.

Scoville, W. B., & Milner, B. (1957). Loss of recent memory after bilateral hippocampal lesions. J Neurol Neurosurg Psychiatry, 20, 11–21.

Squire, L. R. (1992). Memory and the hippocampus: a synthesis from findings with rats, monkeys, and humans. Psychol Rev, 99, 195–231.

Sreenivasan, K. K., & D’Esposito, M. (2019). The what, where and how of delay activity. Nature Reviews Neuroscience, 20, 466–481.

Suh, J., Rivest, A. J., Nakashiba, T., Tominaga, T., & Tonegawa, S. (2011). Entorhinal cortex layer III input to the hippocampus is crucial for temporal association memory. Science, 334, 1415–1420.

Uluc, I., Schmidt, T. T., Wu, Y. H., & Blankenburg, F. (2018). Content-specific codes of parametric auditory working memory in humans. Neuroimage, 183, 254–262.

Umbach, G., Kantak, P., Jacobs, J., Kahana, M., Pfeiffer, B. E., Sperling, M., & Lega, B. (2020). Time cells in the human hippocampus and entorhinal cortex support episodic memory. 2020.2002.2003.932749.

Wolff, M. J., Jochim, J., Akyurek, E. G., & Stokes, M. G. (2017). Dynamic hidden states underlying working-memory-guided behavior. Nat Neurosci, 20, 864–871.

Wolff, M. J., Kandemir, G., Stokes, M. G., & Akyurek, E. G. (2020). Unimodal and Bimodal Access to Sensory Working Memories by Auditory and Visual Impulses. J Neurosci, 40, 671–681.

Zatorre, R. J., Evans, A. C., & Meyer, E. (1994). Neural mechanisms underlying melodic perception and memory for pitch. J Neurosci, 14, 1908–1919.

Zumer, J.M., Brookes, M.J., Stevenson, C.M., Francis, S.T.,Morris, P.G (2010). Relating BOLD fMRI and neural oscialltions through convolution and optimal linear weighting. Neuroimage,49, 1479–1489.

